# The calcium channel Orai1 is required for osteoblast development: studies in a chimeric mouse with variable *in vivo* Runx-cre deletion of Orai-1

**DOI:** 10.1101/2022.02.14.480443

**Authors:** Lisa J Robinson, Jonathan Soboloff, Irina L Tourkova, Quitterie C Larrouture, Dionysios J Papachristou, Scott Gross, Robert Hooper, Elsie Samakai, Paul F Worley, Jan Tuckermann, Michelle R Witt, Harry C Blair

## Abstract

The calcium-selective ion channel Orai1 has a complex role in bone homeostasis, with defects in both bone production and resorption detected in Orai1 germline knock-out mice. To determine whether Orai1 has a direct, cell-intrinsic role in osteoblast differentiation and function, we bred Orai1 flox/flox (Orai1^f/f^) mice with Runx2-cre mice to eliminate its expression in osteoprogenitor cells. Interestingly, Orai1 was expressed in a mosaic pattern in Orai1^f/f^-Runx2-cre bone. Specifically, antibody labeling for Orai1 in vertebral sections was uniform in wild type animals, but patchy regions in Orai1^f/f^-Runx2-cre bone revealed Orai1 loss while in other areas expression persisted. Nevertheless, by micro-CT, bones from Orai1^f/f^-Runx2-cre mice showed reduced bone mass overall, with impaired bone formation identified by dynamic histomorphometry. Cortical surfaces of Orai1^f/f^-Runx2-cre vertebrae however exhibited patchy defects. In cell culture, Orai1-negative osteoblasts showed profound reductions in store-operated Ca^2+^ entry, exhibited decreased alkaline phosphatase activity, and had markedly impaired substrate mineralization. We conclude that defective bone formation observed in the absence of Orai1 reflects an intrinsic role for Orai1 in differentiating osteoblasts.

## Introduction

Orai1 is a calcium-selective ion channel found in the plasma membrane, activated by STIM1, a calcium sensor located on the endoplasmic reticulum [1, 2, 3]. Although expressed in all cells, the extent to which different cell types require Orai1 is variable. For example, T cell activation requires Orai1; its loss causes severe combined immunodeficiency [4, 5]. Loss of Orai1 also causes a wide range of changes in cardiovascular function [6], neuronal function [7, 8], tooth formation [9, 10] and others [11]. We [12, 13, 14, 15, 16] and others [17, 18], demonstrated that Orai1 contributes to bone differentiation and maintenance in conventional knockout mice and *in vitro* cell lines.

Investigation of mice with global deletion of Orai1 revealed loss of multinucleated osteoclasts, that, intriguingly, did not lead to osteopetrosis: Instead, micro-computed tomography showed reduced cortical ossification and thinned trabeculae in Orai1^-/-^animals compared with controls [12]. We attributed the unexpected finding of reduced osteoclast formation without increased bone density to a probable defect in bone formation by osteoblasts. However, considering the near-lethal phenotype of Orai1^-/-^mice [11] and the interdependence of osteoclast and osteoblast differentiation, these investigations were insufficient to draw definitive conclusions regarding the role of Orai1 within bone-forming osteoblasts specifically.

Here, we assessed the role of Orai1 on osteoblast differentiation *in vivo* using a conditional knockout mouse. Interestingly, we found that Orai1 deletion *in vivo* was highly regional, creating a mosaic effect within bone. Irrespective, loss of Orai1 eliminated store-operated Ca^2+^ entry in osteoblasts and precursors resulted in a severe defect in osteoblast differentiation and function. Altogether, this investigation shows that Orai1 is required for normal osteoblast function.

## Materials and Methods

### Animals and Genotyping

Orai1^fl/fl^ mice were generated by flanking Orai1 exons 2/3 with loxP as previously described [16, 19]. Mice were backcrossed to C57Bl/6 for a minimum of 10 generations and then crossed with Runx2-cre mice (Tg(Runx2-icre)1Jtuc) [20] for conditional deletion in the osteoblast lineage. Orai1^fl/fl^ mice were identified by PCR of genomic DNA from tail snips using the following primers: flox-F: ACC CAT GTG GTG GAA AGA AA and flox-R: TGC AGG CAC TAA AGA CGA TG; these generate a 746 bp product in wild-type mice and a 505 bp product in Orai1^fl/fl^ mice [16]. To confirm excision of Orai1 from bone cells, PCR of genomic DNA was performed using the flox-F primer paired with excision-R: CAG AAA GAA CTA CAC AGA GAA ATC, as described [16]; excision results in a 520 bp product, while none is produced when the gene is intact. Mice were sacrificed at 16 weeks unless otherwise noted. For dynamic histomorphometry, animals were injected with xylenol orange (80 µg/g mouse weight) five days prior to sacrifice, then with calcein (20 µg/g mouse weight) 2 days later. Mouse long bones were dissected, cleaned of soft tissue, and processed for isolation of mesenchymal stem cells and osteoblasts. The vertebral column was removed for histologic and microCT analysis. Work was approved by the Temple University IACUC.

### Cell isolation and differentiation

Unless stated, media and chemicals were from Thermo-Fisher. Mesenchymal stem cells (MSC) were isolated from long bones as described [21]. Briefly, marrow was flushed with Minimum Essential Medium-alpha (alphaMEM) containing 10% fetal bovine serum (Sigma-Aldrich, St. Louis, MO), penicillin and streptomycin. After plating cells 16 hours to allow adhesion of stromal cells, non-adherent cells were removed, and the adherent cells grown in selective medium (Mesencult, StemCell Technologies, Cambridge, MA), at 2 × 10^6^ cells/ cm^2^, for proliferation of MSC. Osteoblasts were isolated from long bones as described [22]: briefly, after removal of marrow, the long bones, cut into 1-2 mm fragments, were incubated in 2 mg/ml collagenase type II (260 U/mg, Worthington, Lakewood, NJ) in Dulbecco’s Modified Eagle’s Medium (DMEM) for 2 hours at 37 °C. The bone fragments were then rinsed and transferred to culture dishes with Osteoblast Growth Medium (DMEM with 1 g/l glucose, 10% heat inactivated fetal bovine serum, 30 µg/ml ascorbate, penicillin, streptomycin, and amphotericin B. For differentiation to osteoblasts with mineralized matrix formation, MSC or osteoblasts at confluence were transferred to Osteoblast Mineralization Medium (Osteoblast Growth medium with 30 µg/ml ascorbate and 10 mM 2-glycerol phosphate). During culture, media were replaced every 2-3 days.

### Histomorphometry and Histology

Static histomorphometry was performed as described [16]. Briefly, lumbar vertebrae were fixed overnight in 3.7% formalin, then transferred to 70% ethanol for micro-CT analysis using Bruker Skyscan 1272 with a bone density cutoff of 150 mg/cm^2^, at 5 µm resolution. Scans were analyzed using Bruker CTan software for trabecular parameters; three-dimensional images were produced with Bruker CTvox software. For dynamic histomorphometry, vertebral sections were taken from animals labeled with xylenol orange and calcein two days apart. Lumbar vertebrae 1-3 were used for dynamic histomorphometry and histologic studies, and vertebrae 4-6 were for micro-computed tomography. Bone samples for fluorescent microscopy were cut without decalcification. For hematoxylin and eosin staining, bone was fixed, dehydrated, paraffin embedded, and cut as 10 µm thick sections using a rotary microtome [23]. Cortical thickness was measured orthogonal to the vertical axis in microns at 300 µm intervals; dynamic histomorphometry was as described using calcein and xylenol orange labels [23]. Labeling was analyzed by observers blinded to genotype.

### Alkaline phosphatase, mineralization, and adipocyte assays

Mineral was labeled with silver nitrate (von Kossa stain): cultures were rinsed with water and fixed with 3.7% formalin for 2 minutes, then incubated with 2% AgNO_3_ under UV light for 10 minutes, and rinsed again with water. Alkaline phosphatase activity was determined in cell cultures using 0.01% naphthol phosphate substrate in citrate-buffered saline at pH 8 plus 0.25 mg/ml of fast blue to precipitate an insoluble blue adduct [24]. Parallel cultures were assessed for adipocyte formation by incubating fixed cells in 0.3% Oil red O in 60% isopropanol for 1 hour and rinsed with water.

### Measurement of Store Operated Calcium Entry

Osteoblasts on glass coverslips were loaded with 2 µM Fura-2 AM-ester (Invitrogen) for 30 minutes at 25 °C in 107 mM NaCl, 7.2 mM KCl, 1.2 mM MgCl_2_, 11.5 mM glucose, 20 mM HEPES, 1 mM CaCl_2_, at pH 7.2. Washed cells were allowed to de-esterify dye for 30 minutes at 25°C as described [16]. Ca^2+^ measurements used a Leica DMI 6000B fluorescence microscope controlled by Slidebook Software (Intelligent Imaging; Denver, CO). Intracellular Ca^2+^ is shown as 340/380 nm ratios from single cells. Data used 15-20 cells per mouse and three or more experiments.

### Orai1 labeling

Rabbit polyclonal-anti Orai1 was used for tissue labeling [25]; rabbit anti-Orai1 (extracellular) antibody was from Alomone Labs (ACC-062, Jerusalem, Israel). Triplicate sections from three wild type and three knockout animals (to allow no-antibody controls) of wild type (WT) and knockout bone (KO) were de-paraffinized by 67 °C heating for two days followed by a xylene rinse 12 hours. Labeling was done twice with different sections. In each case, sections were hydrated with 70% ethanol followed by PBS with 1% bovine serum albumin (BSA) with 2 mM EDTA (blocking solution) overnight at room temperature, followed by antibody at 1:100 in PBS with BSA and EDTA (or no antibody controls) overnight at room temperature, and then secondary labeling with anti-rabbit Alexafluor 488 (green) for four hours at room temperature, followed by rinsing and fixation with 3.7% formalin in PBS. When indicated, post staining with phalloidin-rhodamine (1:100) was used to label actin. For Western blots, cell lysates were made using 0.3% SDS, 50 mM tris, pH 7, with proteinase, and phosphatase inhibitors. Proteins were separated on 4-12% gradient gel and transferred to polyvinylidene difluoride (PVDF) membranes. The primary antibody was the Alomone antibody used for fluorescence (1:200); secondary antibody was horse radish peroxidase conjugated anti-rabbit (1:40,000, Jackson ImmunoResearch, West Grove, PA). Proteins on blots were detected by enhanced chemiluminescence detection (ECL plus, Amersham, Piscataway, NJ, USA). Mouse beta-actin antibody (1:1,000, Sigma) with secondary horse radish peroxidase conjugated anti-mouse (1:40,000) were controls.

### Statistics

Results are mean ± SD, for three or more measurements or as stated. Comparisons of differences used Student’s unpaired t-test. Significance indicates p < 0.05.

## Results

### Phenotype of wild type and Orai1^f/f^-Runx2-cre animals

Orai1^f/f^-Runx2-cre conditional knockout mice revealed no gross differences in size, health or behavior compared to controls. Histo-morphometric studies of vertebrae were performed for detailed assessment of the bone. Wild type vertebrae had typical morphology (examples are shown, Fig 1A), while Orai1^f/f^-Runx2-cre vertebrae showed a patchy reduction in cortical bone (examples are shown, Fig 1B). To better understand the basis for this distinct pock-marked appearance in Orai1^f/f^-Runx2-cre animals, cortical bone was examined in cross section. The appearance of the wild type cortex was unremarkable, but cortical bone in Orai1^f/f^-Runx2-cre animals showed irregular thinning (Fig 2A-B). This lack of regularity is unusual but was observed in three of three Orai1^f/f^-Runx2-cre and none of three wild type animals analyzed. Analysis of measurements of cortical thickness, in microns at 300 µm intervals, showed that cortical bone was significantly thinned in the Orai1^f/f^-Runx2-cre animals (Fig 2C). We hypothesized that areas of reduced bone reflected regions in which Orai1 had been deleted by cre-recombination thus impairing bone formation, but that cre-mediated excision was incomplete with areas of normal thickness reflecting preserved bone formation by osteons with Orai1 positive cells. This was further investigated by antibody labeling for Orai1 (see below).

**Fig 1.**
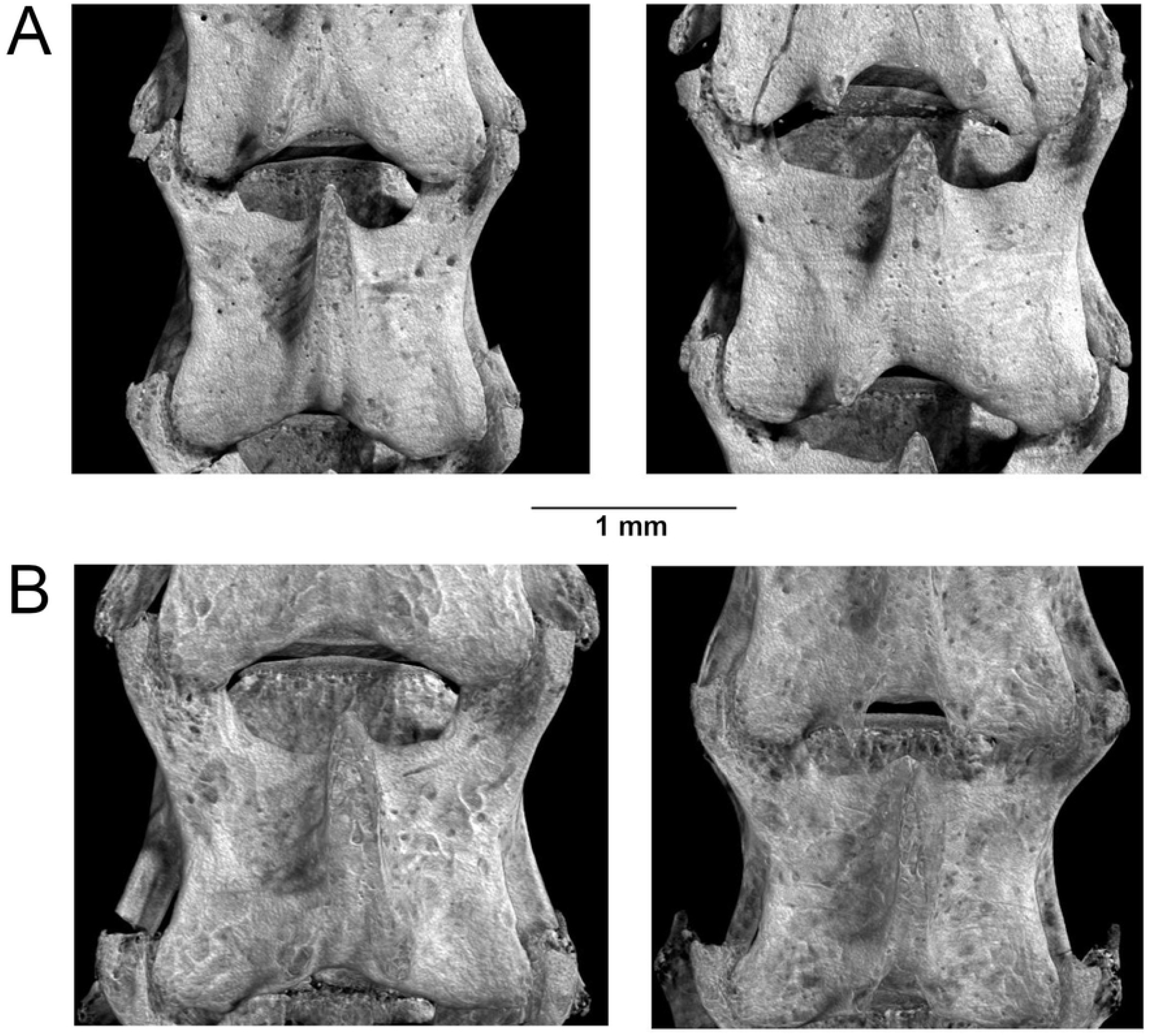
Surface of wild type and Orai1fl/fl-Runx2cre fourth lumbar vertebrae. Bruker CTvox software-generated three-dimensional images of vertebrae reconstructed from microCT scans at 5 µm resolution. All animals were homozygous for floxed Orai1; the conditional knockouts (lower panels) are Runx2-cre positive. **A**. Representative vertebrae from control animals. Apart from sites of blood vessel entry, the surface of the bone is smooth, typical for mice at four months of age. **B**. Representative vertebrae from Orai1fl/fl-Runx2cre animals. In contrast to the control vertebrae, the bone surface appears irregular with patchy darker areas representing regions of reduced bone.

**Fig 2.**
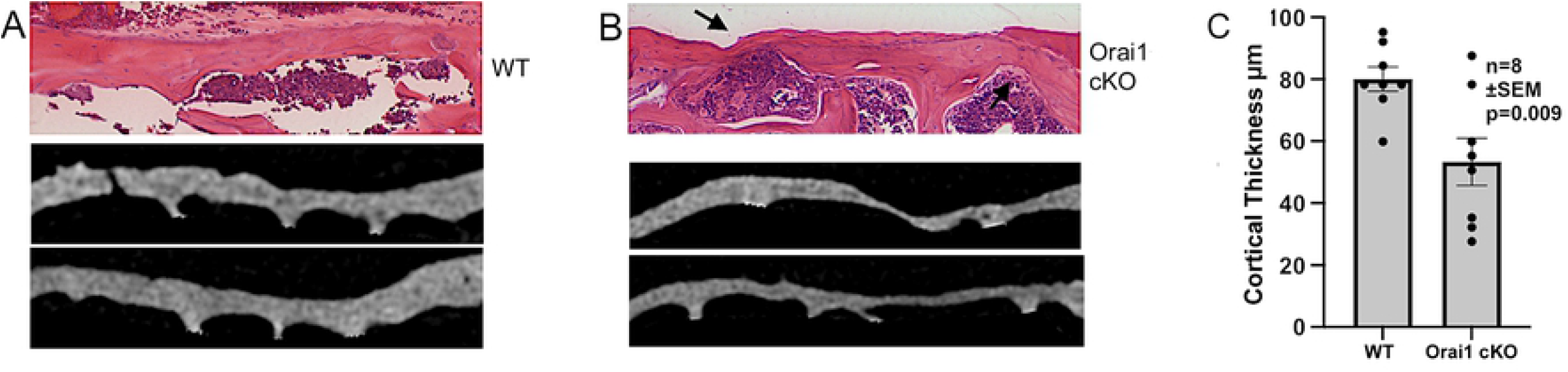
Cross sections of vertebral cortex from wild-type (A) and Orai1^f/f^-Runx2-cre conditional KO (B) animals. Images of H&E stained histologic sections from upper lumbar vertebrae (top panels) are 1 mm wide; images from microCT scans of lower lumbar vertebrae (middle and lower panels) are 1.4 mm wide, with trabecular bone deleted. **A**. Wild type cortex with typical smooth bone showed relatively uniform thickness. **B**. The Orai1fl/fl-Runx2cre (abbreviated Orai1 cKO) animals had irregularly thinned regions (arrow) but no other distinguishing features. This is in keeping with the appearance of the bone surface in the three-dimensional reconstructions (see Fig 1). **C**. Cortical bone thickness, though variable, is reduced on average in Orai1fl/fl-Runx2cre (Orai1 cKO) animals (p = 0.009, N=8).

### WT and Orai1^f/f^-Runx2-cre animals show differences by static and dynamic histomorphometry

Standard histomorphometric parameters for trabecular bone were determined by microCT. Results revealed that overall bone mass, measured as bone volume/tissue volume (BV/TV), was reduced in Orai1^f/f^-Runx2-cre animals compared to controls (Fig 3A; p=0.01). Mean trabecular thickness was also decreased (Fig 3B; p<0.01) in Orai1^f/f^-Runx2-cre bone, as was trabecular number (Fig 3C, p<0.02) in the Orai1 conditional knockouts. Antemortem fluorescent labeling of newly formed bone by injected calcein and xylenol orange was used for dynamic histomorphometry studies. Consistent with the patchy thinner and thicker areas of bone noted above (Figs 1 and 2) suspected to reflect regions of Orai1 negative and positive osteoblasts, we found a patchy distribution of new bone formation with calcein/xylenol orange labeling (Fig 3D) in the Orai1 conditional knock-out animals. This irregular labeling does not fit traditional histomorphometric parameters and was found in none of the controls, but was consistent across the Orai1^f/f^-Runx2-cre samples examined. In regions where labeling was intact, dynamic histomorphometric measurements at time of sacrifice showed no difference in bone apposition rate (interlabel distance, not shown) or fraction of labeled bone surface (Fig 3E). Finally, despite the irregularity of the phenotype, the calcein labeling did demonstrate reduction in bone deposition overall in Orai1^f/f^-Runx2-cre mice (Fig 3F, p = 0.03). This indicates that bone forming regions in Orai1^f/f^-Runx2-cre mice were smaller than in control animals. Given the patchy nature of these effects, we suspect that the contribution of Orai1 to bone formation in affected regions may be variable. To determine if indeed variable Orai1 deletion is the cause of this variable phenotype, we assessed Orai1 deletion further.

**Fig 3.**
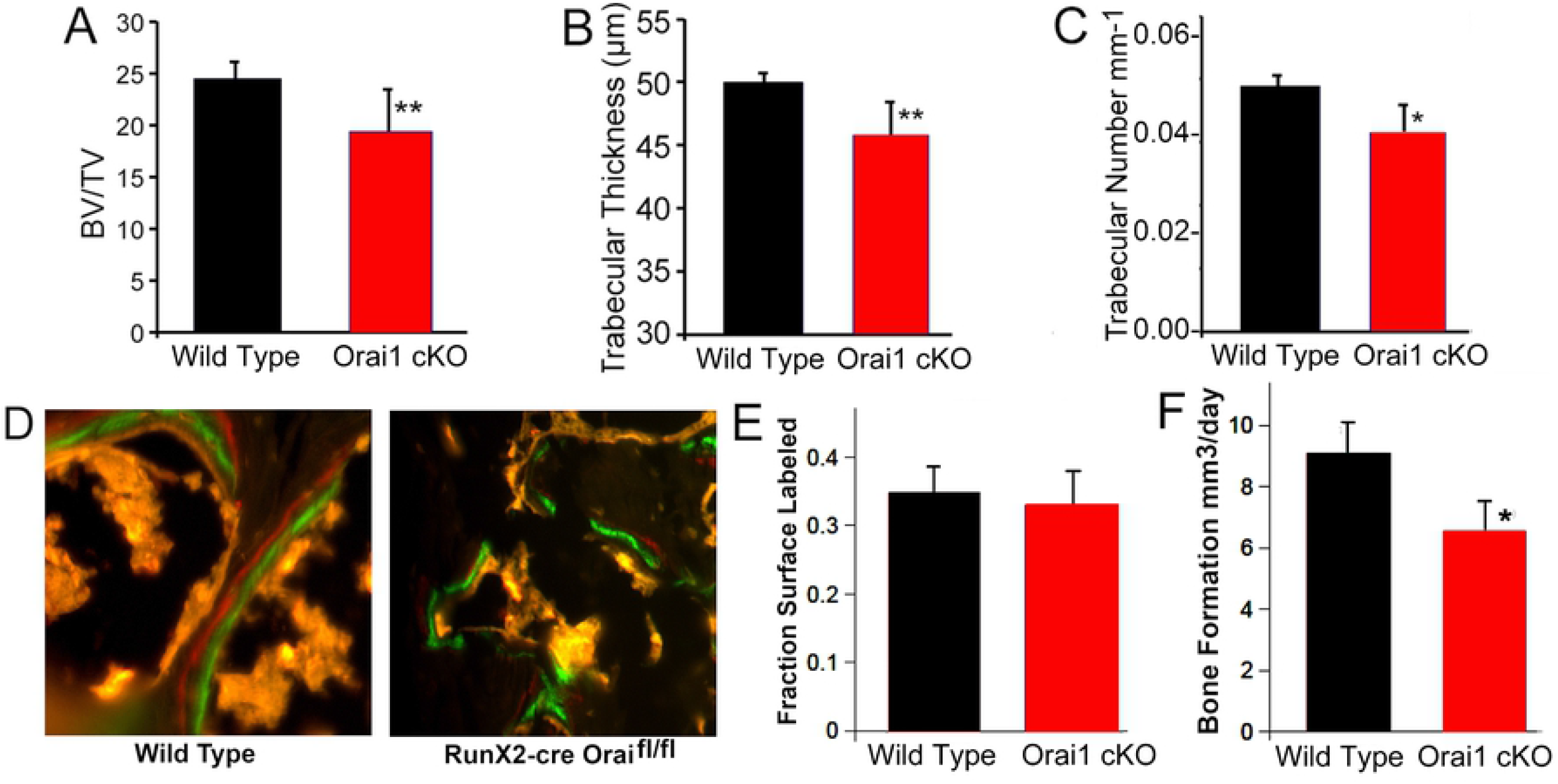
Key features of WT and conditional KO animals on static and dynamic histomorphometry. **A**. Bone volume/total volume (BV/TV) of vertebrae. Vertebral bone is significantly reduced (N=7, p=0.01) in the Orai1fl/fl-Runx2cre (Orai1 cKO) mice compared to controls. The greater variability in the conditional-knockout bone is thought to reflect variable cre excision; this variability is also apparent in the cortical section thickness (Fig 2; see text). **B**. Trabecular thickness is significantly decreased in the Orai1 conditional knockout mice (N=7, p<0.01). **C**. Trabecular number is also significantly decreased in the conditional knockouts (N=7, p<0.02). **D**. Example of xylenol orange (red) and calcein (green) fluorescent labels in wild type and conditional KO bones. The interlabel distance was not statistically different (not illustrated). Fields shown are 100 µm wide. **E**. The proportion of calcein labeled bone surface in wild type and Orai1fl/fl-Runx2cre bone (abbreviated Orai1 cKO) was not statistically different (N=4). **F**. The bone formation rate at 4 months of age was significantly reduced in the Orai1fl/fl-Runx2cre mice (Orai1 cKO) compared to controls (N=4, p=0.03).

### Variable deletion of osteoblast flox/flox Orai1 by the Runx2-cre promoter

Incomplete deletion by Cre has been demonstrated in many different mouse models (see Discussion), although we found the apparent mosaic phenotype of this particular mouse striking. To demonstrate variable deletion of Orai1 within bone cells, we performed antibody labeling using an Orai1 antibody in wild type and Orai1^f/f^-Runx2-cre bone (Fig 4). Controls for specificity of labeling are shown (Fig 4A). While all WT bone cells expressed Orai1 (Fig 4A), there were patches of Orai1-negative (arrows) as well as Orai1-positive osteocytes in Orai1^f/f^-Runx2-cre mice (Fig 4B and 4C, lower panels). Western blot showed significant but incomplete Orai1 deletion in cultures of Orai1^f/f^-Runx2-cre osteoblasts (Fig 4D). This was consistent variable deletion of Orai1 in osteoblasts from Orai1^f/f^-Runx2-cre mice. To determine the functional implications of Orai1 deletion in osteoblasts, *in vitro* cell cultures, negative for Orai1 by PCR, in mineralization medium were used, as described below.

**Fig 4.**
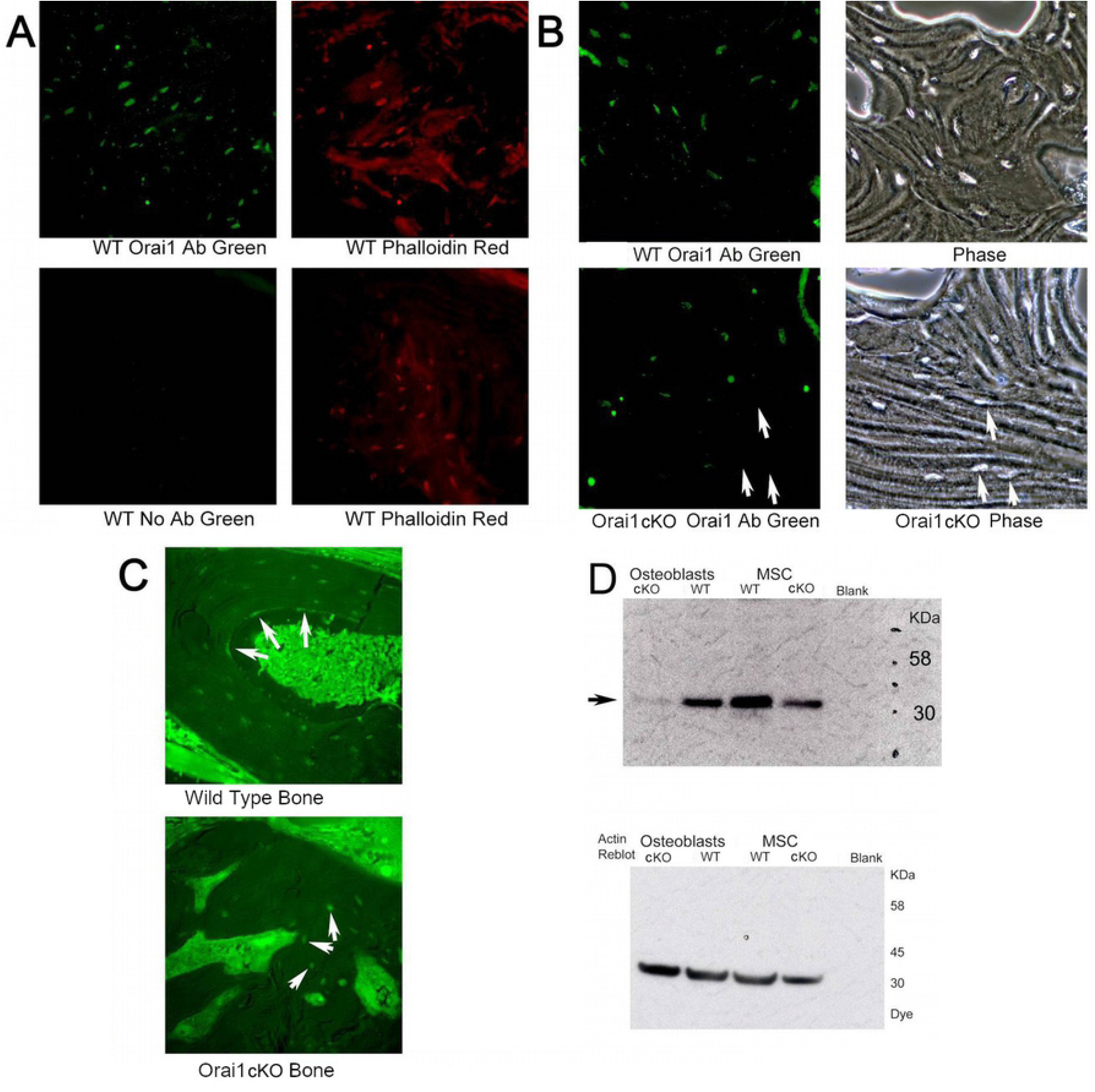
Variable efficacy of Runx2-cre in deleting flox/flox Orai1 in osteoblasts. It is often assumed that promoter-cre constructs uniformly delete the target in all cells of an organ. Recent findings suggest that for reasons that are not clear this is sometimes not the case (see Text), we tested this by antibody labeling of Orai1 in wild type and Orai1fl/fl-Runx2cre bone. **A**. Rabbit antibody detects Orai1 in osteoblasts (upper left panel). Absence of antibody eliminates labeling (lower left panel). All sections are from one wild type animal. Left and right panels are of the same section. Osteoblasts are shown independently with phalloidin rhodamine (right panels). Fields are 200 µm across. **B**. In the wild type animal, osteoblasts are shown with the antibody at high power (fields 200 microns wide) and in phase of the same field (right). In the lower panels, the same field of a conditional KO animal. Some of the conditional KO cells (Orai1fl/fl-Runx2cre) do not label (arrows, phase and antibody label, lower panels). **C**. Lower power fields, 350 µm, of Orai1 labeled osteoblasts (surrounding tissue fluorescence is an artifact) in wild type bone (top) and conditional KO bone (bottom). Osteoblasts in the wild type label strongly, including surface cells of the bone (arrows, top). Some, but not all, of the osteoblasts in the conditional KO label (arrows, bottom). **D**. Western blot for Orai1 in cells expressing or not expressing the protein (See Fig 5). Thirty-five µg loads of cell protein from the isolates indicated were run on denaturing SDS-PAGE and blotted. This blot using the very specific Alomone antibody shows a trace of Orai1 even in the osteoblast KO preparation (left lane) and reduced, but not absent, Orai1 in the MSC KO (upper panel). The beta actin re-blot (lower panel) confirms similar protein loads.

### Effect of Orai1 deletion on store-operated calcium entry

MSCs were collected from Orai1fl/fl and Orai1^f/f^-Runx2-cre mice before differentiating into osteoblasts *in vitro* and measuring store-operated calcium entry as described [16]. Cells were treated with thapsigargin in the absence of extracellular calcium to deplete the endoplasmic reticulum of calcium and initiate the store-operated calcium response. While a typical store-operated calcium response was observed in Orai1-expressing osteoblasts, loss of Orai1 completely eliminated store-operated calcium entry (Fig 5). These observations show that Orai1 is required for store-operated calcium entry in osteoblasts.

**Fig 5.**
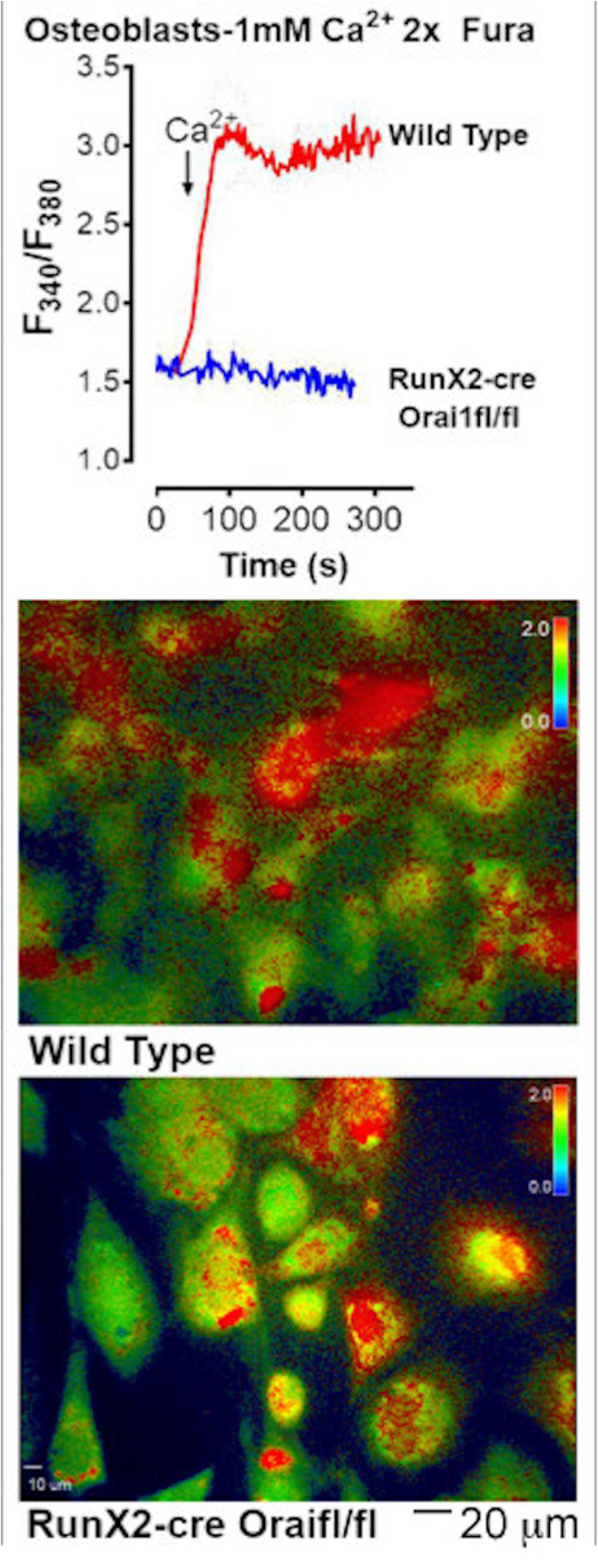
Calcium activated calcium release in cells from wild type and Runx2cre-Orai^fl/fl^ mice. Intracellular calcium was measured by Fura2 (Wang, 2009) (top panel). Cells were treated with thapsigargin in the absence of extracellular calcium. Only the wild type cells show calcium entry upon extracellular calcium repletion. Examples of Fura2 signals from control and conditional KO cells are shown in the middle and lower panels respectively, with false color reflecting calcium concentration in cells (scale is shown in upper right corner of each panel). A scale bar for size for the photomicrographs is shown below the bottom panel.

### Orai1^f/f^-Runx2-cre cells showed severe defects in osteoblast differentiation

We analyzed osteoblast differentiation using wild type cells and cells from conditional knockouts in which absence of Orai1 was confirmed by PCR (not illustrated), grown in mineralization medium with 2-glycerol phosphate and ascorbate. Mineral deposition was detected using silver nitrate (von Kossa); mineralization was prominent in cultures of wild type cells, but markedly reduced for Orai1^f/f^-Runx2-cre osteoblasts (Fig 6A). Additionally, alkaline phosphatase positivity in the conditional KO was significantly decreased relative to wild type (Fig 6B). Reduced osteoblastic differentiation correlates with increased adipocytes in mice fed a Western diet [26]. However, no Orai1-dependent differences in adipogenic differentiation were identified by Oil red O staining in these culture conditions (Fig 6C), suggesting that Orai1 expression was did not favor osteoblast-adipocyte conversion. For definitive evaluation of the role of Orai1 further studies using adipogenic culture conditions would be needed.

**Fig 6.**
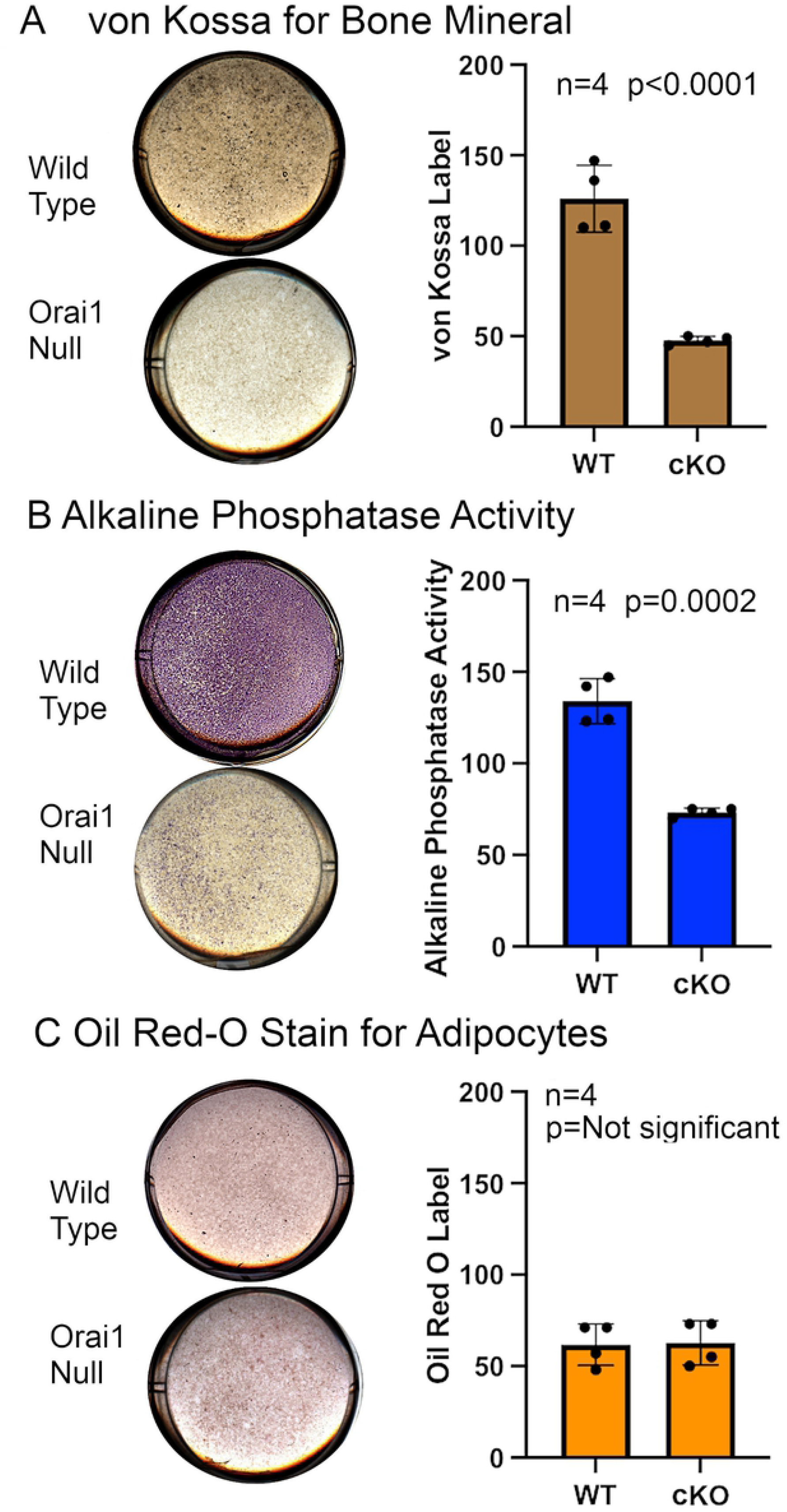
Elimination of Orai1 results in profoundly reduces OB differentiation and mineralization from OB precursors. Osteoblasts isolated as described in the methods section from control or Runx2-cre floxed (Orai1^f/f^-Runx2-cre) conditional knock-out animals, were incubated 24 days in osteoblast mineralization medium and analyzed by histologic staining. Each well illustrated is 2 cm across. **A**. Von Kossa staining for mineral. Wild type cells made mineral nodules, but there were only rare and small nodules in Orai1 knockout cell cultures. Representative culture wells for control (WT) and conditional knockout (cKO) cells are shown on the left. Staining was quantified for four samples of each genotype. Mineralized matrix production appeared significantly reduced in cultures of Orai1-deficient osteoblasts (p<0.0001, N=4). **B**. Alkaline phosphatase activity. Representative cultures are shown on the left. Results quantified from four samples of each genotype are shown on the right. Wild type cells produced much more alkaline phosphatase, with small nodules of high activity as in the silver stain for mineral (A), there was uniformly less alkaline phosphatase in Orai1 null cells (p=0.0002, N=4). **C**. Oil red O staining for adipocytes. Representative cultures are shown on the left. Staining was quantified for four samples of each genotype with results shown on the right. There was no evidence of increased adipogenic differentiation in Orai1 null cells, with minimal labeling in cultures of either genotype.

## Discussion

We show that Orai1 serves an essential cell-intrinsic function in osteoblast differentiation. Interpretation is complicated by incomplete deletion of Orai1, resulting in animals exhibiting an intriguing mosaic phenotype. Interestingly, complete deletion of glucocorticoid genes early in osteoblast differentiation has been reported in Runx2-cre mice [20]. While the reasons for this difference are not clear; epigenetic differences in the status of the Orai1 gene amongst different osteoblasts represent one possible explanation; silencing of genes due to histone methylation have been shown to interfere with cre recombinases in prior study [27]. While this possibility has not been established for the Runx2-cre floxed Orai1 combination, it would be consistent with the findings reported here and a subject for future study. In keeping, other studies have shown incomplete cre-mediated excision [28, 29].

Global analysis of bone density showed significant but subtle differences. We feel that this likely reflects the incomplete deletion of Orai1 *in vivo*. However, *in vitro* bone differentiation from Orai1^-/-^cells caused profound reductions in osteoblast differentiation. This reflects that cells used were either wild type or confirmed Orai1^-/-^, while *in vivo* a patchwork of normal and Orai1^-/-^cells occurs.

Further, while Runx2 is crucial for the commitment of mesenchymal stem cells to the osteoblast lineage, Runx2 expression is down-regulated at later stages [30]. While there is convincing evidence that Orai1 is involved in osteoblast differentiation and function in the current study and in [31], the uncertain status of Orai1 in osteoblast precursors combined with the temporary nature of Runx2 expression *in vivo* may contribute to the mosaic phenotype in this model.

The present work using non-transformed cells indicates unequivocally the essential role of Orai1 in bone differentiation. When Orai1 is eliminated in pre-osteoblastic cells, limited osteoblast differentiation occurs, with remarkable variability of osteoblast-related differentiation. The ability of osteoblasts to form mineralizing nodules is completely abrogated in Orai1-null osteoblasts. This is consistent with the necessity of store-operated calcium entry for bone differentiation. This further suggests that the formation of advanced osteoblast-related structures [32] is impossible without store-operated calcium entry. There are a many studies indicating importance for calcium signals in bone formation, e.g., [33, 34, 35], but previously not showing specificity for Orai1 or calcium-activated calcium release in normal osteoblast differentiation. Our work is cell specific, unlike general Orai1 knockout studies [12, 31], in which bone degrading, bone forming, and potentially other cell types were affected. In truly Orai1 negative osteoblasts, bone differentiation is severely compromised.

## Acknowledgements

We thank Cayla R Sudano and Kelechi M Onwuka for assistance.

